# Flanker: a tool for comparative genomics of gene flanking regions

**DOI:** 10.1101/2021.02.22.432255

**Authors:** William Matlock, Samuel Lipworth, Bede Constantinides, Timothy E.A. Peto, A. Sarah Walker, Derrick Crook, Susan Hopkins, Liam P. Shaw, Nicole Stoesser

**Author notes:** Joint first authors (William Matlock, Samuel Lipworth). Joint senior authors (Liam P. Shaw, Nicole Stoesser).

## Abstract

Analysing the flanking sequences surrounding genes of interest is often highly relevant to understanding the role of mobile genetic elements (MGEs) in horizontal gene transfer, particular for antimicrobial resistance genes. Here, we present Flanker, a Python package which performs alignment-free clustering of gene flanking sequences in a consistent format, allowing investigation of MGEs without prior knowledge of their structure. These clusters, known as ‘flank patterns’, are based on Mash distances, allowing for easy comparison of similarity across sequences. Additionally, Flanker can be flexibly parameterised to finetune outputs by characterising upstream and downstream regions separately and investigating variable lengths of flanking sequence. We apply Flanker to two recent datasets describing plasmid-associated carriage of important carbapenemase genes (blaOXA-48 and blaKPC-2/3) and show that it successfully identifies distinct clusters of flank patterns, including both known and previously uncharacterised structural variants. For example, Flanker identified four Tn4401 profiles that could not be sufficiently characterised using TETyper or MobileElementFinder, demonstrating the utility of Flanker for flanking gene characterisation. Similarly, using a large (n=226) European isolate dataset, we confirm findings from a previous smaller study demonstrating association between Tn1999.2 and *bla_OXA-48_* upregulation and demonstrate 17 flank patterns (compared to the 5 previously identified). More generally the demonstration in this study that flank patterns are associated with to geographical regions and antibiotic susceptibility phenotypes suggests that they may be useful as epidemiological markers. Flanker is freely available under an MIT license at https://github.com/wtmatlock/flanker.

**Data Summary:** NCBI accession numbers for all sequencing data used in this study is provided in Supplementary Table 1. The analysis performed in this manuscript can be reproduced in a binder environment provided on the Flanker Github page (https://github.com/wtmatlock/flanker).

## Introduction

The increasing incidence antimicrobial resistance (AMR) in clinical isolates poses a threat to all areas of medicine [1–3]. AMR genes (ARGs) are found in a diverse range of genetic contexts, bacterial species, and in both clinical and non-clinical environments (e.g., agricultural, refuse and natural ecosystems) [4–7]. However, the mechanisms underpinning the dissemination of many ARGs between these reservoirs remain poorly understood, limiting the efficacy of surveillance and the ability to design effective interventions. Usually, ARGs are spread vertically, either via chromosomal integration or stable association of a plasmid within a clonal lineage, or by horizontal gene transfer (HGT) through mobile genetic elements (MGEs) e.g., transposons or plasmids [8]. HGT can accelerate the rate of ARG acquisition, both within and across species [9–11].

The epidemiology of ARGs can therefore involve multiple levels, from clonal spread to MGEs. There are many existing software tools to facilitate epidemiological study of bacterial strains [12–16], whole plasmids [17, 18], and smaller MGEs [19, 20]. Several tools and databases exist for the annotation of non-plasmid MGEs such as insertion sequences (ISs) and transposons [19, 20], but all rely on comparisons to reference sequences, so are limited to known diversity. Reference-free tools for analysing MGE diversity would therefore be a useful addition. Here we describe a Flanker, a simple, reference-free tool to investigate MGEs by analysing the flanking sequences of ARGs.

The flanking sequences (hereafter, ‘flanks’) around an ARG that has been mobilised horizontally may act as signatures of relevant MGEs and support epidemiological analyses. However, these flanks can contain a great deal of structural variation due to their evolutionary history. Where a single known MGE is under investigation, it is possible to specifically type this element (for example, using TETyper [19]) or align flanks against a known ancestral form after the removal of later structural variation [21]. However, often multiple structures may be involved. This is particularly true for ARGs which move frequently on a variety of MGEs. Studies of different ARGs often choose different *ad hoc* approaches to extract flanks and cluster genetic structures. Examples include hierarchical clustering of isolates carrying an ARG based on short-read coverage of known ARG-carrying contigs [22], assigning assembled contigs into ‘clustering groups’ based on gene presence and synteny [23] or iterative ‘splitting’ of flanks based on pairwise nucleotide BLAST identity [24]. A consistent and simple approach for this task would not only avoid repeated method development, but also aid comparison between methods developed for specific ARGs.

To address this problem, we developed Flanker, a pipeline to analyse the regions around a given ARG in a consistent manner. Flanker flexibly extracts the flanks of a specified gene from a dataset of contigs, then clusters these sequences using Mash distances to identify consistent structures [25]. Flanker is available as a documented Python and Bioconda package released under the MIT open-source license. Source code is deposited at https://github.com/wtmatlock/flanker and documentation at https://flanker.readthedocs.io/en/latest/.

## Methods

### Flanker

The Flanker package contains two basic modules: the first extracts a region of length *w* around an annotated gene of interest, and the second clusters such regions based on a user-defined Mash distance threshold (default --threshold 0.001, Fig. 1a). Within each FASTA/multi-FASTA format input file, the location of the gene of interest is first determined using the Abricate annotation tool [31]. Flanks around the gene (optionally including the gene itself to enable complete alignments with --include_gene) are then extracted and written to a FASTA format file using BioPython [39]. Flanker gives users the option to either extract flanks using a single window (defined by length in base-pairs [bp]) or multiple windows from a start position (--window) to an end position (--wstop) in fixed increments (--wstep). Flanks may be extracted from upstream, downstream or on both sides of the gene of interest (--flank). Corrections are also made for circularised genomes where the gene occurs close to the beginning or end of the sequence (--circ mode) and for genes found on both positive and negative strands. The clustering module groups flanks of user-defined sequence lengths together based on a user-defined Mash [25] distance threshold (--threshold) of user-defined sequence lengths.

**Figure 1:**
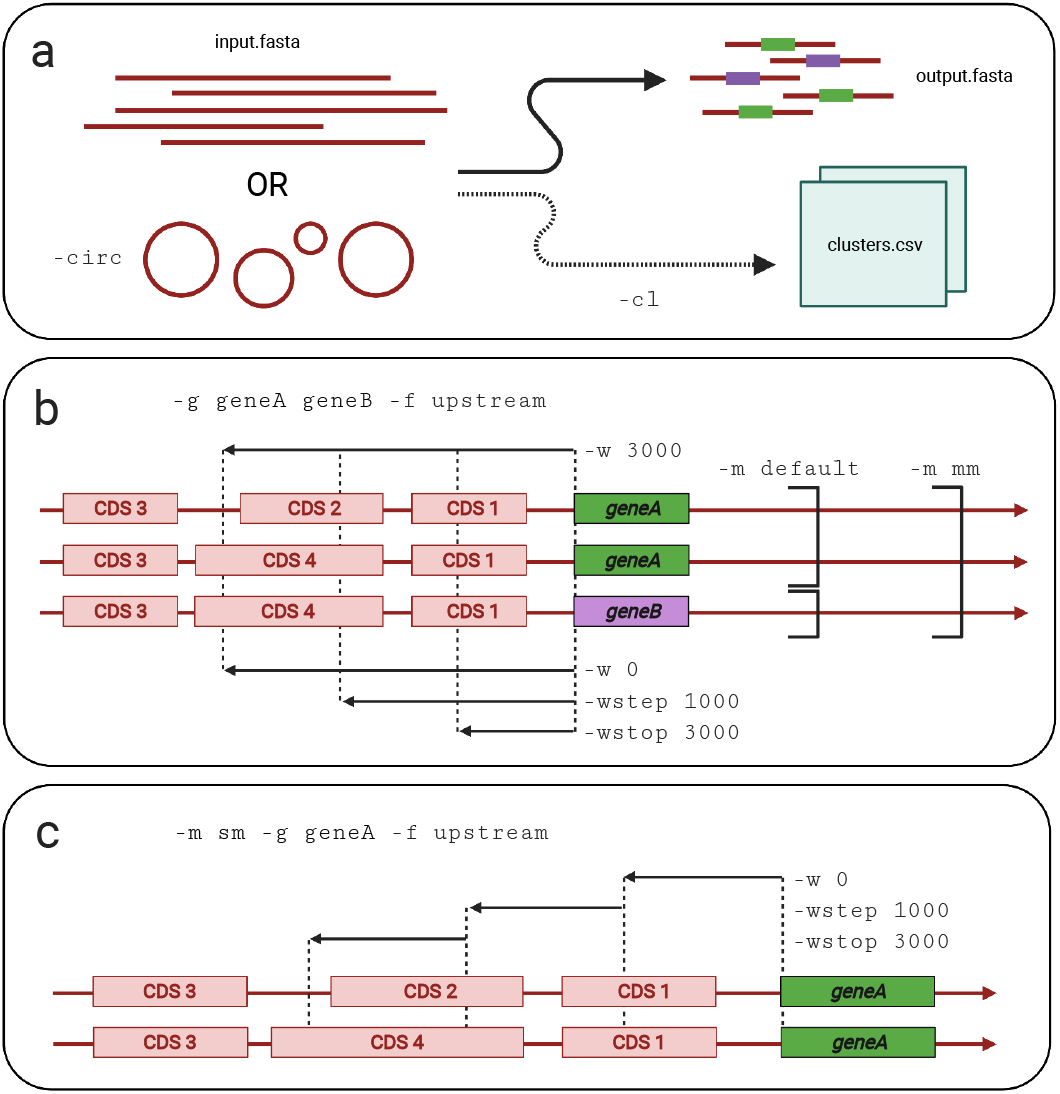
Schematic of Flanker’s modes and parameters. (a) Flanker uses Abricate to annotate the gene of interest in input sequences and outputs associated flanking sequences, optionally clustering (-cl) these on a user defined Mash distance threshold. It can take linear or circularised sequences. (b) In this example, genes *geneA* and *geneB* have been queried (-g geneA geneB), and only the upstream flank is desired (-f upstream). The top single black arrow represents choosing a single window of length 3000bp (-w 3000), whereas the bottom three black arrows represent stepping in 1000bp windows from 0bp to 3000bp (-w 0 -wstep 1000 -wstop 3000). The default mode (-m default) extracts flanks for all annotated alleles separately, but the multi-allelic mode (-m mm) extracts flanks for all alleles in parallel. (c) Flanker has a supplementary salami mode (-m sm), which outputs non-contiguous blocks of sequence with a start point, step size, and end point (-w 0 -wstep 1000 -wstop 3000), represented by the three black arrows.

In default mode (--mode default), Flanker considers multiple gene queries in turn. In multi-allelic mode (--mode mm), Flanker considers all genes in the list for each window (Fig. 1b). Multiple genes can be queried by either a space-delimited list in the command line (--gene geneA geneB), or a newline-delimited file with the list of genes option (--list_of_genes). A supplementary module ‘salami mode’ (--mode sm) is provided to allow comparison of non-contiguous blocks from a start point (--window), step size (--wstep) and end point (--wstop) (Fig. 1c).

### Datasets

To validate Flanker, demonstrate its application and provide a comparison with existing tools, we used two recent datasets of complete plasmids (derived from hybrid long-/short-read assemblies) containing carbapenemase genes of clinical importance [23, 26]. The first dataset comprised 51 complete *bla*_OXA-48_-harbouring plasmids; 42/51 came from carbapenem-resistant *Escherichia coli* and *Klebsiella pneumoniae* isolates from patients in the Netherlands [26] and 9/51 from EuSCAPE (a European surveillance programme investigating carbapenem resistance in Enterobacterales) [23]. The second dataset comprised 50 *bla*_KPC-2_ or *bla*_KPC-3_-(*Klebsiella pneumoniae* carbapenemase)-harbouring plasmids in carbapenem-resistant *K. pneumoniae* isolated from the Netherlands [26] (8/50) and as part of the EuSCAPE study (42/50)[23]. The EuSCAPE dataset [23, 27] additionally contains a large collection of short-read sequencing data for *Klebsiella* spp. isolates alongside meropenem susceptibility data. This was used to demonstrate additional possible epidemiological applications of the Flanker tool by evaluating whether specific flank patterns were more likely to be associated with phenotypic meropenem resistance.

### Mash distances

Pair-wise distances between flanks were calculated using Mash (version 2.2.2) [25]. Mash reduces sequences to a fixed-length MinHash sketch, which is used to estimate the Jaccard distance between k-mer content. It also gives the Mash distance, which ranges from 0 (~identical sequences) to 1 (~completely dissimilar sequences). We used the default Mash parameters in all analyses. The Mash distance was developed to approximate the rate of sequence mutation between genomes under a simple evolutionary model, and explicitly does not model more complex processes. We use it here for fast alignment-free clustering of sequences and do not draw any direct conclusions about evolution from pairwise comparisons.

### Clustering

To cluster the flanks, Flanker generates an adjacency matrix weighted by Mash distances. It then thresholds this matrix to retain edges weighted less than or equal to the defined threshold. This is then used to construct a graph using the Python NetworkX library [28] and clusters are defined using the nx.connected_components function, which is analogous to single linkage. This is a similar methodology to that used by the Assembly De-replicator tool [29] (from which Flanker re-uses several functions). However, Flanker aims to assign all flanks to a cluster rather than to deduplicate by cluster.

### Cluster validation

We validated the output of flanking sequence-based clustering using a PERMANOVA test, implemented with the adonis function from the Vegan package (version 2.7.5) [30] in R. Only flanks in clusters of at least two members were considered; 42/51 and 48/50 of *bla*_OXA-48_ and *bla*_KPC-2/3_ flanks, respectively. The formula used was Mash dist ~ cluster, with the ‘euclidean’ method and 999 permutations.

### Comparison to existing methods/application

We compared the classifications of TETyper (v1.1) [19] and MEFinder (v1.0.3) [20] to those produced by Flanker for 500bp and 5000bp flanks around *bla*_KPC-2/3_ genes. TETyper was run using the-threads 8 and --assemblies options with the *Tn4401* reference and SNP/structural profiles provided in the package and MEFinder was run in Abricate[31] using the –mincov 10 option. For comparisons of the proportions of resistant isolates per flank pattern (denoted FP), isolates were classified as resistant or sensitive using the European Committee on Antimicrobial Susceptibility Testing (EUCAST) breakpoint for meropenem (>8mg/L) [32].

### Data visualisation

All figures were made using Biorender (https://biorender.com) and the R packages ggplot2 (v3.3.0) [33], gggenes (v0.4.0) [34] and ggtree (v2.4.1) [35] Prokka (v1.14.6) [36] was used to annotate Flanker output. Mashtree (v1.2.0) [37] was used to construct a visual representation of Mash distances between whole plasmid genomes. Plasmidfinder was used to detect the presence/absence of plasmid types using Abricate (version 1.01) with --mincov 80 and --minid 80 [38]. Galileo AMR (https://galileoamr.arcbio.com/mara/) was used to visualise the transposon variants. Figures can be reproduced using the code in the GitHub repository (https://github.com/wtmatlock/flanker).

## Results

### Clustering validation and comparison with TETyper/MEFinder

The clustering mode was validated numerically with a PERMANOVA test (Mash dist ~ cluster: *bla*_OXA-48_ *p*-value < 0.001, *bla*_KPC2/3_ *p*-value < 0.001; see Methods). Figures 2 and 3 also provide a visual comparison of an alignment of genes (‘Gene graphical representation’ panel) to the Flank pattern.

**Figure 2:**
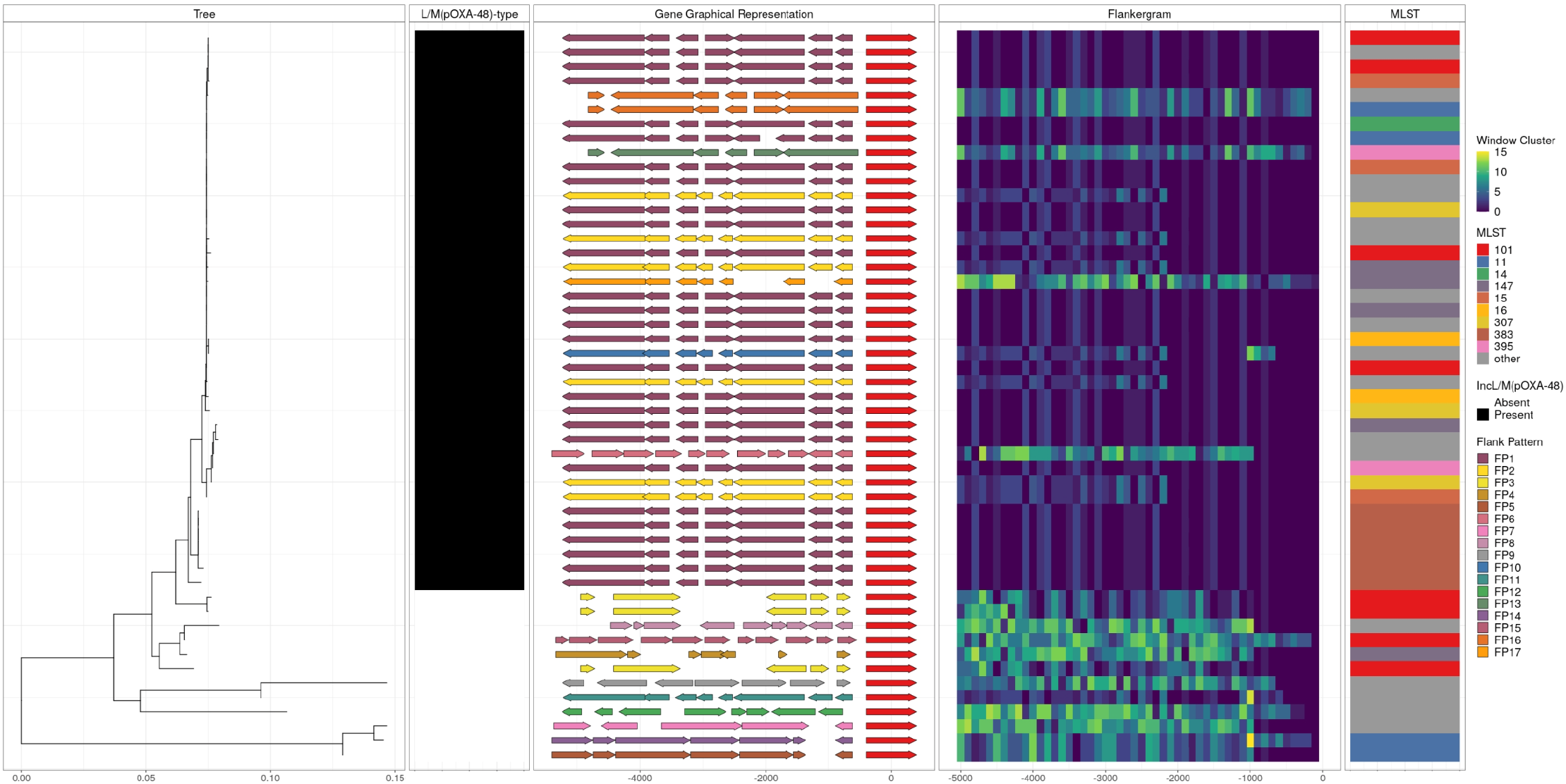
Flanking regions 5000bp upstream of *bla*_OXA-48_ carrying plasmids from *Klebsiella* pneumoniae isolates. The ‘Tree’ panel is a Neighbour-Joining tree constructed from Mash distances between complete sequences of plasmids carrying the *bla*_OXA-48_ gene. The second panel indicates the presence/absence of a L/M(pOXA-48)-type plasmid. The ‘Gene Graphical Representation’ panel schematically represents coding regions in the 5000bp sequence upstream of the *bla*_OXA-48_ gene, which is shown in red. Other genes are coloured according to the flank pattern which considers the overall pattern of all 100bp window clusters up to 2200bp (the approximate upstream limit of Tn*1999*). The “Flankergram” shows window clusters of all groups over each 100bp window between 0 and 5000bp. The dotted line at 2200bp indicates the approximate point of upstream divergence between several flank patterns. The ‘MLST’ panel shows *K. pneumoniae* multi-locus sequence types (MLSTs), with those occurring once labelled ‘other’. FPs are numbered in ascending order according to abundance in the hybrid assemblies. Data used to make this figure came from the Dutch CPE surveillance and EUSCAPE hybrid assembly datasets.

**Figure 3:**
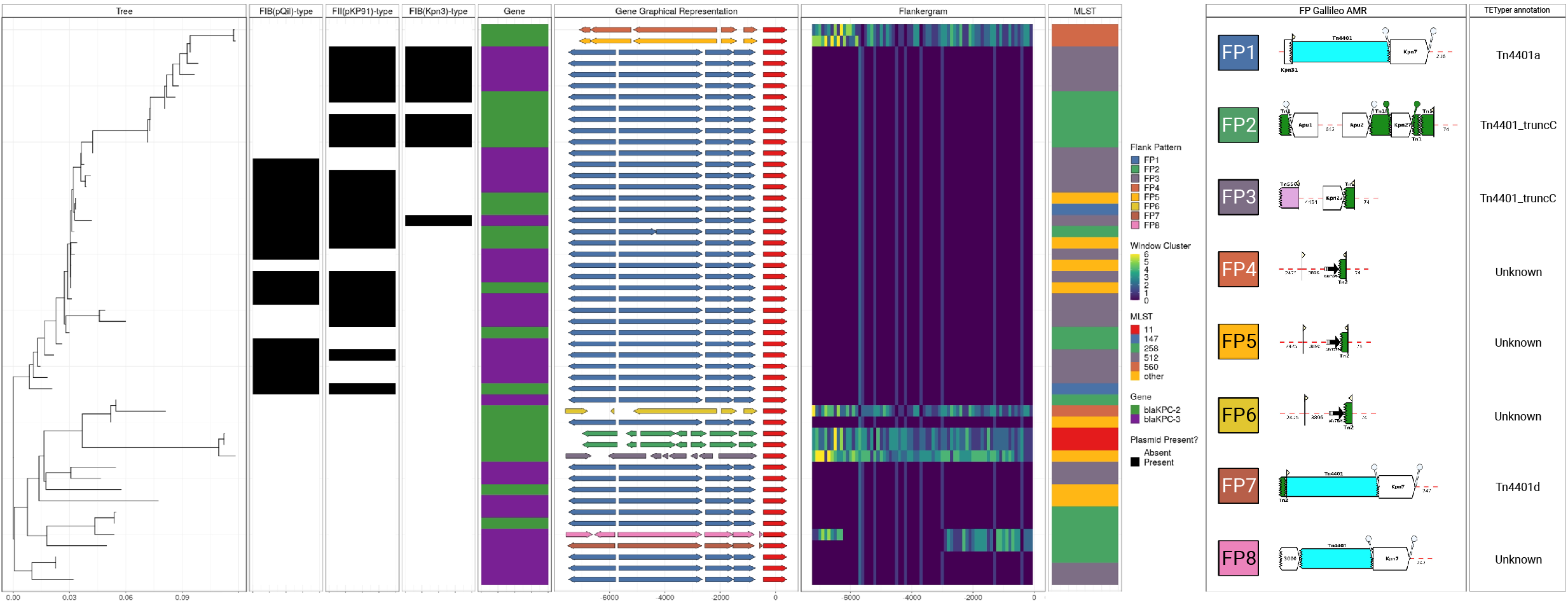
Flanking regions 7200bp upstream of *bla*_KPC-2/3_ carrying plasmids from *Klebsiella pneumoniae* isolates. The ‘Tree’ panel is a Neighbour-Joining tree constructed from Mash distances between complete sequences of plasmids carrying the *bla*_KPC-2/3_ gene. The next three panels indicate the presence/absence of FIB(pQ1I)-, FII(pKP91)-, and FIB(Kpn3)-type plasmids. The ‘Gene’ column indicates which *bla*_KPC_ allele (2 or 3) is present. The ‘Gene Graphical Representation’ panel schematically represents coding regions in the 7200bp sequence region upstream of the *bla*_KPC-2/3_ gene, which is shown in red. Other genes are coloured according to the flank pattern, which here takes into account the overall pattern of all 100bp window groups (shown in the “Flankergram” panel) over the full 7200bp region upstream of *bla_KPC-2/3_*. The “Flankergram” shows window clusters over each 100bp window between 0 and 7200bp. The ‘MLST’ panels shows *K. pneumoniae* MLSTs, with those occurring once labelled ‘other’. The final two panels show the Galileo AMR and the TETyper outputs for the 8 FPs, respectively. The FPs are numbered in ascending order according to abundance in the hybrid assemblies.

Of the two existing tools we compared in evaluating the flanks around *bla*_KPC2/3_, TETyper was by far the slowest (1172 seconds [s]), whereas MEFinder, run in Abricate, and Flanker took 7s and 11s, respectively (benchmarked on 5000bp upstream flanks on a cluster with Intel Skylake 2.6GHz chips). MEFinder was able to detect *Tn4401* but could not provide any further structural resolution and was unable to classify 6/50 (12%) 500bp and 1/50 (2%) 5000bp flanks. TETyper structural profiles were consistent with Flanker when analysing 500 and 5000bp upstream regions (Figure 3), though Flanker split a group of six isolates with the TETyper structural profile 1-7127|7202-10006 into four groups (Table S2). To map our FPs to the established nomenclature, we additionally compared the output of Flanker to that of TETyper when the latter was given the entire *Tn4401* sequence (i.e., by evaluating the typical 7,200bp Tn*4401*-associated flank upstream of *bla_KPC_*). Flanker and TETyper classifications of Tn*4401* regions were broadly consistent (Table S2), though this analysis demonstrated the potential benefit of the reference-free approach of Flanker which showed that four non-Tn*4401* structural profiles (‘unknown’ in TETyper) were distinct from each other. In addition, TETyper classified 3 flanks as Tn*4401*_truncC-1, whereas Flanker resolved this cluster into two distinct groups (Table S2).

### Application to plasmids carrying *bla*_OXA-48_

The carbapenemase gene *bla*_OXA-48_ has been shown to be disseminated by Tn*1999*-associated structures (~5kb, see detailed review in [40]) nested in L/M-type plasmids, and as part of an IS1R-associated composite transposon containing *bla*_OXA-48_ and part of Tn*1999*, namely *Tn6237* (~21.9kb), that has been implicated in the chromosomal integration of *bla*_OXA-48_ [27, 41]. It has been recently demonstrated that most *bla*_OXA-48_-like genes in clinical isolates in Europe are carried on highly similar L/M(pOXA-48)-type plasmids, with evidence of both horizontal and vertical transmission across a diverse set of sequence types [23]. Whilst Tn*1999*-like flanking regions are relatively well characterised [40], in this example we chose an initial arbitrary upstream window of 5000bp to simulate a scenario in which there is no prior knowledge. Inspection of a plot of window clusters (i.e., as shown in the ‘Flankergram’ in Fig. 2) demonstrates that Flanker output allows the empirical identification of the position ~2200bp upstream of *bla*_OXA-48_ as an important point of structural divergence without requiring annotation (as shown at ~2200 along the *x*-axis, where the window cluster colour schemes diverge), corresponding to the edge of Tn*1999* at its expected position.

Using complete plasmids from the Netherlands [26]/EUSCAPE [23] hybrid assembly datasets, Flanker identified 17 distinct FPs in the 2200bp upstream sequence of *bla*_OXA-48_ of which seven occurred in L/M(pOXA-48)-type plasmids (Fig. 2, Table S3). To investigate the association of phenotypic carbapenem resistance with *bla*_OXA-48_ FPs, we created a Mash sketch using one randomly chosen representative per group and screened an Illumina sequenced collection of European carbapenemase-resistant *Klebsiella* isolates [27] for containment (n=425) (Mash screen, assigning FP based on the top hit [median identity = 1.00; range: 0.97-1.00]). Two FPs (FP6 and FP16) accounted for 338/425 (80%) of isolates; both were widely distributed across Europe. Of the 226 isolates with meropenem susceptibility data available, those belonging to FP6 were proportionally more meropenem-resistant compared to FP16 (70/135 [52%] vs. 6/44 [14%], exact *p*-value<0.001; Fig.3). Annotation (using Galileo AMR; see methods) of these revealed that whereas FP16 contains Tn*1999*,FP6 contains *Tn1999.2*, which has been previously described as creating a strong promoter which produces 2-fold higher enzymatic activity [42].

### Application to plasmids carrying *bla*_KPC-2/3_

David et al. showed that *bla*_KPC-2/3_ genes have been disseminated in European *K. pneumoniae* clinical isolates via a diverse collection of plasmids in association with a dominant clonal lineage, ST258/512, which accounted for 230/312 (74%) of *bla*_KPC_-associated isolates in the EuSCAPE collection [23]. *bla*_KPC_ has largely been associated with variants of a ~10kb transposon, Tn*4401* [43, 44]. From the combined EuSCAPE [23] and Dutch CPE collection [26] of 50 hybrid assembled KPC-containing plasmids, Flanker identified 8 distinct FPs over a 7200bp window upstream of bla_KPC-2/3_ (Fig. 4; Table S2). This window length was chosen to capture the entire Tn*4401* sequence upstream of *bla_KPC_*.

Considering Mash containment of the 8 representative FPs within the EuSCAPE short read assemblies dataset, 346/442 (78%) belonged to FP1 (corresponding to isoform Tn*4401*a). Whilst FP1 was widely distributed across Europe, FP2 (corresponding to *Tn4401*_truncC) and FP7 (corresponding to Tn*4401*d) appeared more geographically restricted: FP2 to Spain (5/5, 100%) and FP7 to Israel (19/59, 32%) and Portugal (34/59, 58%) with isolates also found in Poland and Germany (n=2 each) and Italy and Austria (n=1, Table S3). Of the 442 short read assemblies, 274 had meropenem MIC data available for analysis. There was no evidence of a difference in the proportion of isolates resistant to meropenem between FP1 and FP7 (202/238 [85%] vs 23/25 [92%], exact *p*-value=0.5, Table S4), though there was incomplete susceptibility data for isolates from both groups (108/346 [31%] for FP1 and 38/63 [60%] for FP7).

## Discussion

We present Flanker, a fast and flexible Python package for analysing gene flanking sequences. We anticipate that this kind of analysis will become more common as the number of complete reference-grade, bacterial assemblies increase. Our analysis of data from the EuSCAPE project suggests that flank patterns (FPs) might be useful epidemiological markers when evaluating geographical associations of sequences. Additionally, we validated findings of a small (n=7) PCR-based study on a large (n=226) European dataset, confirming an association between Tn1999.2 and increased meropenem resistance. A key advantage compared to existing tools is that there is no reliance on reference sequences or prior knowledge. Despite analysing only a relatively small number (n=50) of complete *bla_KPC_* containing plasmids, there were four distinct FPs which TETyper classified as ‘unknown’ because their profiles had not been previously characterised. Similarly, we identified 17 FPs associated with *bla*_OXA-48_ in contrast to the five structural variants of *Tn1999* currently described in the literature.

TETyper works well when alleles/structural variants are known but can only classify a single transposon type at a time and requires manual curation when this is not the case. The observed diversity of flanking sequences is likely to continue to increase and manual curation of naming schemes will be arduous to maintain. MEFinder on the other hand is a quick screening tool which can search a large library of known mobile elements but lacks sequence level resolution. Whilst Flanker overcomes these challenges, users may need to perform downstream analysis to interpret its output. We hope that Flanker will be complementary to these and other similar existing tools by reducing the dimensionality of large datasets and identifying smaller groups of sequences to focus on in detail. Though we have developed Flanker for ARGs, Abricate allows use of custom databases meaning any desired genes of interest could be analysed. Accurate outputs from Flanker will be dependent on the quality of input assemblies, and on the correct annotation of the gene of interest.

In summary, we present Flanker, a tool for comparative genomics of gene flanking regions which integrates several existing tools (Abricate, Biopython, NetworkX) in a convenient package with a simple command-line interface.

## Authors and contributors

Contributions have been attributed by the CRediT system as follows:

Conceptualisation: WM, SL, LPS, NS

Methodology: WM, SL

Software: WM, SL, BC

Validation: WM, SL

Formal Analysis: WM, SL

Investigation: WM, SL

Resources: DWC, TP, ASW, NS

Data Curation: SL, WM

Writing - Original Draft Preparation: SL, WM, LPS, NS

Writing - Review and Editing: SL, WM, LS, BC, NS, DC, TP, ASW

Visualisation: SL, WM

Supervision: LPS, NS, TP, ASW, DC

Project Administration: SL, WM, NS, LPS

Funding: TP, DC, ASW, NS

## Data availability

Accessions for the plasmid sequences, and MEFinder and TETyper outputs are provided in Table S1.

## Acknowledgements

The authors thank the EuSCAPE and Dutch CPE surveillance groups for making their data publicly available.

## Impact statement

The global dissemination of antimicrobial resistance genes (ARGs) has in part been driven by carriage on mobile genetic elements (MGEs) such as transposons and plasmids. However, our understanding of these MGEs remains poor, partly due to their high diversity. This means current referenced based approaches are often inappropriate. ‘Flanker’ is a fast software tool which overcomes this barrier by *de* novo clustering of ARG flank diversity by sequence similarity. We demonstrate the utility of Flanker by associating *bla_OXA-48_* and *bla_KPC-2/3_* flanking sequences with geographic regions and resistance phenotypes.

