## Supplement for "Flanker: a tool for comparative genomics of gene flanking regions"

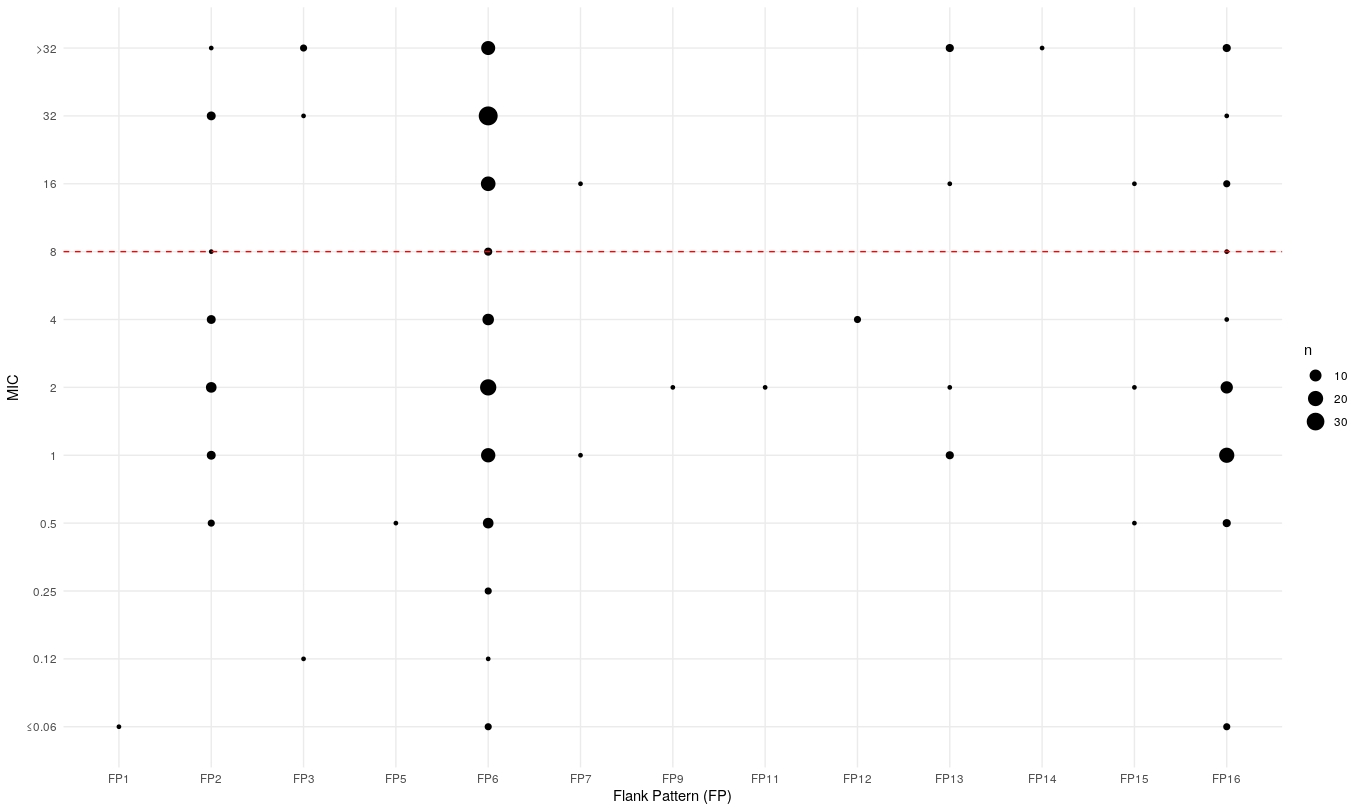

**Figure S1**: Distribution of meropenem minimum inhibitory concentrations (MIC mg/L) by Flank Pattern (FP). Circle size indicates the number of observations per category and the red hashed line denotes the EUCAST breakpoint (>8mg/L). Isolates from the EuSCAPE dataset (short read only sequencing, n=226 isolates with linkable phenotype data) were assigned to a Flank Pattern based on the top mash containment hit). No isolates were assigned to FPs 4,8 or 17 and the one isolate assigned to FP10 had no linkable phenotypic data.

**Table S1** – NCBI project accession numbers for sequencing data used

| Description | Project Accession | Number of isolates |
| --- | --- | --- |
| Complete plasmid assemblies containing *bla_KPC2/3_* | PRJEB33308 (EuSCAPE)  ERP118777 (Dutch CPE) | 42  8 |
| Complete plasmid assemblies containing *bla_OXA-48_* | PRJEB33308 (Dutch CPE)  PRJNA591727 (EuSCAPE) | 9  42 |
| Short read Illumina fastq files for isolates containing *bla_OXA-48*_* | PRJEB10018 (EuSCAPE) | 425 |
| Short read Illumina fastq files for isolates containing *bla_KPC-2/3*_* | PRJEB10018 (EuSCAPE) | 442 |

*Full list of accession numbers used and associated metadata is available at <https://doi.org/10.6084/m9.figshare.14074250.v1>

**Table S2** – Flank Pattern (FP), MEFinder and TETyper calls for blaKPC-2/3 flanking regions. The isolate column gives the contig name within the assembly, the next three columns give FP assignations at 500/5000/7200bp upstream of the gene. Columns 5 to 8 give the MEFinder/TETyper calls for the 500bp/5000bp upstream flanking regions whereas the final column gives TETyper calls when TETyper was given the whole plasmid assembly as input.

| **Contig Name** | **Flanker 500bp** | **Flanker 5000bp** | **Flanker 7200bp** | **MEFinder 500bp** | **MEFinder 5000bp** | **TETyper 500bp** | **TETyper 5000bp** | **TETyper Tn4401 whole** |
| --- | --- | --- | --- | --- | --- | --- | --- | --- |
| CABFYD010000003.1 | 1 | 1 | 1 | Tn4401\|1\|KT378596.1 | Tn4401\|1\|KT378596.1 | 1-6602\|7020-7118\|7202-10006 | 1-2202\|7020-7118\|7202-10006 | Tn4401a |
| MT560073.1 | 1 | 1 | 1 | Tn4401\|1\|KT378596.1 | Tn4401\|1\|KT378596.1 | 1-6602\|7020-7118\|7202-10006 | 1-2202\|7020-7118\|7202-10006 | Tn4401a |
| CABFYQ010000002.1 | 1 | 1 | 1 | Tn4401\|1\|KT378596.1 | Tn4401\|1\|KT378596.1 | 1-6602\|7020-7118\|7202-10006 | 1-2202\|7020-7118\|7202-10006 | Tn4401a |
| CABFZV010000006.1 | 1 | 1 | 1 | Tn4401\|1\|KT378596.1 | Tn4401\|1\|KT378596.1 | 1-6602\|7020-7118\|7202-10006 | 1-2202\|7020-7118\|7202-10006 | Tn4401a |
| CABGAG010000002.1 | 1 | 1 | 1 | Tn4401\|1\|KT378596.1 | Tn4401\|1\|KT378596.1 | 1-6602\|7020-7118\|7202-10006 | 1-2202\|7020-7118\|7202-10006 | Tn4401a |
| CABFYI010000002.1 | 1 | 1 | 1 | Tn4401\|1\|KT378596.1 | Tn4401\|1\|KT378596.1 | 1-6602\|7020-7118\|7202-10006 | 1-2202\|7020-7118\|7202-10006 | Tn4401a |
| CABGAO010000002.1 | 1 | 1 | 1 | Tn4401\|1\|KT378596.1 | Tn4401\|1\|KT378596.1 | 1-6602\|7020-7118\|7202-10006 | 1-2202\|7020-7118\|7202-10006 | Tn4401a |
| CABFYF010000002.1 | 1 | 1 | 1 | Tn4401\|1\|KT378596.1 | Tn4401\|1\|KT378596.1 | 1-6602\|7020-7118\|7202-10006 | 1-2202\|7020-7118\|7202-10006 | Tn4401a |
| CABFYH010000003.1 | 1 | 1 | 1 | Tn4401\|1\|KT378596.1 | Tn4401\|1\|KT378596.1 | 1-6602\|7020-7118\|7202-10006 | 1-2202\|7020-7118\|7202-10006 | Tn4401a |
| MT560060.1 | 1 | 1 | 1 | Tn4401\|1\|KT378596.1 | Tn4401\|1\|KT378596.1 | 1-6602\|7020-7118\|7202-10006 | 1-2202\|7020-7118\|7202-10006 | Tn4401a |
| CABGAI010000003.1 | 1 | 1 | 1 | Tn4401\|1\|KT378596.1 | Tn4401\|1\|KT378596.1 | 1-6602\|7020-7118\|7202-10006 | 1-2202\|7020-7118\|7202-10006 | Tn4401a |
| CABGAZ010000003.1 | 1 | 1 | 1 | Tn4401\|1\|KT378596.1 | Tn4401\|1\|KT378596.1 | 1-6602\|7020-7118\|7202-10006 | 1-2202\|7020-7118\|7202-10006 | Tn4401a |
| CABFYZ010000002.1 | 1 | 1 | 1 | Tn4401\|1\|KT378596.1 | Tn4401\|1\|KT378596.1 | 1-6602\|7020-7118\|7202-10006 | 1-2202\|7020-7118\|7202-10006 | Tn4401a |
| CABGAQ010000009.1 | 1 | 1 | 1 | Tn4401\|1\|KT378596.1 | Tn4401\|1\|KT378596.1 | 1-6602\|7020-7118\|7202-10006 | 1-2202\|7020-7118\|7202-10006 | Tn4401a |
| CABGBS010000004.1 | 1 | 1 | 1 | Tn4401\|1\|KT378596.1 | Tn4401\|1\|KT378596.1 | 1-6602\|7020-7118\|7202-10006 | 1-2202\|7020-7118\|7202-10006 | Tn4401a |
| CABFYU010000005.1 | 1 | 1 | 1 | Tn4401\|1\|KT378596.1 | Tn4401\|1\|KT378596.1 | 1-6602\|7020-7118\|7202-10006 | 1-2202\|7020-7118\|7202-10006 | Tn4401a |
| CABGAR010000003.1 | 1 | 1 | 1 | Tn4401\|1\|KT378596.1 | Tn4401\|1\|KT378596.1 | 1-6602\|7020-7118\|7202-10006 | 1-2202\|7020-7118\|7202-10006 | Tn4401a |
| MT560075.1 | 1 | 1 | 1 | Tn4401\|1\|KT378596.1 | Tn4401\|1\|KT378596.1 | 1-6602\|7020-7118\|7202-10006 | 1-2202\|7020-7118\|7202-10006 | Tn4401a |
| CABFZA010000002.1 | 1 | 1 | 1 | Tn4401\|1\|KT378596.1 | Tn4401\|1\|KT378596.1 | 1-6602\|7020-7118\|7202-10006 | 1-2202\|7020-7118\|7202-10006 | Tn4401a |
| CABFXY010000002.1 | 1 | 1 | 1 | Tn4401\|1\|KT378596.1 | Tn4401\|1\|KT378596.1 | 1-6602\|7020-7118\|7202-10006 | 1-2202\|7020-7118\|7202-10006 | Tn4401a |
| CABFZG010000005.1 | 1 | 1 | 1 | Tn4401\|1\|KT378596.1 | Tn4401\|1\|KT378596.1 | 1-6602\|7020-7118\|7202-10006 | 1-2202\|7020-7118\|7202-10006 | Tn4401a |
| CABFYK010000005.1 | 1 | 1 | 1 | Tn4401\|1\|KT378596.1 | Tn4401\|1\|KT378596.1 | 1-6602\|7020-7118\|7202-10006 | 1-2202\|7020-7118\|7202-10006 | Tn4401a |
| CABFXS010000003.1 | 1 | 1 | 1 | Tn4401\|1\|KT378596.1 | Tn4401\|1\|KT378596.1 | 1-6602\|7020-7118\|7202-10006 | 1-2202\|7020-7118\|7202-10006 | Tn4401a |
| CABFYY010000002.1 | 1 | 1 | 1 | Tn4401\|1\|KT378596.1 | Tn4401\|1\|KT378596.1 | 1-6602\|7020-7118\|7202-10006 | 1-2202\|7020-7118\|7202-10006 | Tn4401a |
| CABGAY010000002.1 | 1 | 1 | 1 | Tn4401\|1\|KT378596.1 | Tn4401\|1\|KT378596.1 | 1-6602\|7020-7118\|7202-10006 | 1-2202\|7020-7118\|7202-10006 | Tn4401a |
| CABGBH010000002.1 | 1 | 1 | 1 | Tn4401\|1\|KT378596.1 | Tn4401\|1\|KT378596.1 | 1-6602\|7020-7118\|7202-10006 | 1-2202\|7020-7118\|7202-10006 | Tn4401a |
| CABFYC010000004.1 | 1 | 1 | 1 | Tn4401\|1\|KT378596.1 | Tn4401\|1\|KT378596.1 | 1-6602\|7020-7118\|7202-10006 | 1-2202\|7020-7118\|7202-10006 | Tn4401a |
| CABFYE010000002.1 | 1 | 1 | 1 | Tn4401\|1\|KT378596.1 | Tn4401\|1\|KT378596.1 | 1-6602\|7020-7118\|7202-10006 | 1-2202\|7020-7118\|7202-10006 | Tn4401a |
| CABGAK010000004.1 | 1 | 1 | 1 | Tn4401\|1\|KT378596.1 | Tn4401\|1\|KT378596.1 | 1-6602\|7020-7118\|7202-10006 | 1-2202\|7020-7118\|7202-10006 | Tn4401a |
| CABFYG010000003.1 | 1 | 1 | 1 | Tn4401\|1\|KT378596.1 | Tn4401\|1\|KT378596.1 | 1-6602\|7020-7118\|7202-10006 | 1-2202\|7020-7118\|7202-10006 | Tn4401a |
| CABGAX010000009.1 | 1 | 1 | 1 | Tn4401\|1\|KT378596.1 | Tn4401\|1\|KT378596.1 | 1-6602\|7020-7118\|7202-10006 | 1-2203\|7020-7118\|7202-10006 | Tn4401a |
| CABFXR010000002.1 | 1 | 1 | 1 | Tn4401\|1\|KT378596.1 | Tn4401\|1\|KT378596.1 | 1-6602\|7020-7118\|7202-10006 | 1-2202\|7020-7118\|7202-10006 | Tn4401a |
| CABFYB010000005.1 | 1 | 1 | 1 | Tn4401\|1\|KT378596.1 | Tn4401\|1\|KT378596.1 | 1-6602\|7020-7118\|7202-10006 | 1-2202\|7020-7118\|7202-10006 | Tn4401a |
| CABFYS010000003.1 | 1 | 1 | 1 | Tn4401\|1\|KT378596.1 | Tn4401\|1\|KT378596.1 | 1-6602\|7020-7118\|7202-10006 | 1-2202\|7020-7118\|7202-10006 | Tn4401a |
| CABFYL010000004.1 | 1 | 1 | 1 | Tn4401\|1\|KT378596.1 | Tn4401\|1\|KT378596.1 | 1-6602\|7020-7118\|7202-10006 | 1-2202\|7020-7118\|7202-10006 | Tn4401a |
| CABFYT010000008.1 | 1 | 1 | 1 | Tn4401\|1\|KT378596.1 | Tn4401\|1\|KT378596.1 | 1-6602\|7020-7118\|7202-10006 | 1-2202\|7020-7118\|7202-10006 | Tn4401a |
| MT560078.1 | 1 | 1 | 1 | Tn4401\|1\|KT378596.1 | Tn4401\|1\|KT378596.1 | 1-6602\|7020-7118\|7202-10006 | 1-2202\|7020-7118\|7202-10006 | Tn4401a |
| CABGBM010000006.1 | 1 | 1 | 1 | Tn4401\|1\|KT378596.1 | Tn4401\|1\|KT378596.1 | 1-6602\|7020-7118\|7202-10006 | 1-2202\|7020-7118\|7202-10006 | Tn4401a |
| CABFYN010000003.1 | 1 | 1 | 1 | Tn4401\|1\|KT378596.1 | Tn4401\|1\|KT378596.1 | 1-6602\|7020-7118\|7202-10006 | 1-2202\|7020-7118\|7202-10006 | Tn4401a |
| CABFZZ010000003.1 | 1 | 1 | 1 | Tn4401\|1\|KT378596.1 | Tn4401\|1\|KT378596.1 | 1-6602\|7020-7118\|7202-10006 | 1-2202\|7020-7118\|7202-10006 | Tn4401a |
| CABFZH010000004.1 | 1 | 1 | 1 | Tn4401\|1\|KT378596.1 | Tn4401\|1\|KT378596.1 | 1-6602\|7020-7118\|7202-10006 | 1-2202\|7020-7118\|7202-10006 | Tn4401a |
| CABFYM010000002.1 | 1 | 1 | 1 | Tn4401\|1\|KT378596.1 | Tn4401\|1\|KT378596.1 | 1-6602\|7020-7118\|7202-10006 | 1-2202\|7020-7118\|7202-10006 | Tn4401a |
| MT560080.1 | 2 | 4 | 5 | No hits | Tn1000\|1\|X60200.1 | 1-7127\|7202-10006 | 1-7127\|7202-10006 | unknown |
| MT560061.1 | 2 | 4 | 4 | No hits | Tn1000\|1\|X60200.1 | 1-7127\|7202-10006 | 1-7127\|7202-10006 | unknown |
| MT560063.1 | 2 | 3 | 6 | No hits | Tn1000\|1\|X60200.1 | 1-7127\|7202-10006 | 1-7127\|7202-10006 | unknown |
| CABFYR010000005.1 | 3 | 2 | 2 | No hits | Tn6296\|1\|FJ628167 | 1-7127\|7202-10006 | 1-7127\|7202-10006 | Tn4401_truncC |
| CABGBR010000009.1 | 3 | 2 | 2 | No hits | Tn6296\|1\|FJ628167 | 1-7127\|7202-10006 | 1-7127\|7202-10006 | Tn4401_truncC |
| MT560066.1 | 3 | 5 | 3 | No hits | No hits | 1-7127\|7202-10006 | 1-7127\|7202-10006 | Tn4401_truncC |
| CABFYJ010000003.1 | 4 | 1 | 8 | Tn4401\|1\|KT378596.1 | Tn4401\|1\|KT378596.1 | 1-6633\|7008-7075\|7202-10006 | 1-2233\|7008-7075\|7202-10006 | unknown |
| CABFZC010000002.1 | 4 | 1 | 7 | Tn4401\|1\|KT378596.1 | Tn4401\|1\|KT378596.1 | 1-6633\|7008-7075\|7202-10006 | 1-2233\|7008-7075\|7202-10006 | Tn4401d |

| **Flank Pattern** | **N** |
| --- | --- |
| FP1 | 1 |
| FP2 | 37 |
| FP3 | 8 |
| FP4 | 0 |
| FP5 | 4 |
| FP6 | 230 |
| FP7 | 2 |
| FP8 | 0 |
| FP9 | 5 |
| FP10 | 1 |
| FP11 | 4 |
| FP12 | 2 |
| FP13 | 9 |
| FP14 | 1 |
| FP15 | 13 |
| FP16 | 108 |
| FP17 | 0 |

Table S3 – Number of bla*_OXA-48_* Flank Patterns (FPs) observed in the EuSCAPE short read assembly dataset. For each FP identified by Flanker in the hybrid assembly dataset (2200bp upstream of the gene), one random representative was chosen (see main methods). EuSCAPE short read assemblies (n=425 total) were screened for containment of these using mash and the FP was taken to be the match with greatest containment.

|  | **Group 1** | **Group 2** | **Group 7** | **Group 8** |
| --- | --- | --- | --- | --- |
| Austria | 0 | 0 | 1 | 0 |
| Belgium | 0 | 0 | 0 | 0 |
| Croatia | 0 | 0 | 0 | 0 |
| France | 0 | 0 | 0 | 0 |
| Germany | 0 | 0 | 2 | 1 |
| Greece | 0 | 0 | 0 | 3 |
| Ireland | 0 | 0 | 0 | 0 |
| Israel | 0 | 0 | 19 | 0 |
| Italy | 0 | 0 | 1 | 0 |
| Luxembourg | 0 | 0 | 0 | 0 |
| Macedonia | 0 | 0 | 0 | 0 |
| Poland | 0 | 0 | 2 | 0 |
| Portugal | 0 | 0 | 34 | 0 |
| Romania | 0 | 0 | 0 | 0 |
| Slovakia | 0 | 0 | 0 | 0 |
| Spain | 0 | 5 | 0 | 0 |
| United Kingdom (England, Wales & N. Ireland) | 0 | 0 | 0 | 2 |

**Table S4** – Geographical distribution of bla_KPC-2/3_ flanking patterns observed in the EuSCAPE short read assembly dataset (n=313 with linkable geographic data). For each FP identified by Flanker in the hybrid assembly dataset (7200bp upstream of the gene), one random representative was chosen (see main methods). EuSCAPE short read assemblies were screened for containment of these using mash and the FP was taken to be the match with greatest containment. There were no isolates assigned to FPs3/4/5/6 that had linkable geographic data.

| **Meropenem MIC (mg/L)** | **FP 1** | **FP 2** | **FP 7** | **FP 8** |
| --- | --- | --- | --- | --- |
| ≤0.06 | 0 | 0 | 0 | 0 |
| 0.12 | 0 | 1 | 0 | 0 |
| 0.25 | 0 | 0 | 0 | 0 |
| 0.5 | 0 | 0 | 0 | 0 |
| 1 | 0 | 0 | 0 | 0 |
| 2 | 0 | 1 | 1 | 0 |
| 4 | 0 | 1 | 1 | 0 |
| 8 | 0 | 1 | 0 | 1 |
| 16 | 0 | 0 | 3 | 3 |
| 32 | 0 | 0 | 6 | 1 |
| >32 | 0 | 1 | 14 | 1 |

**Table S5** – Distribution of meropenem minimum inhibitory concentrations (MIC) by blaKPC-2/3 7200bp upstream flanking region in the EuSCAPE dataset (n= 274 with linkable phenotypic data). For each FP identified by Flanker in the hybrid assembly dataset (7200bp upstream of the gene), one random representative was chosen (see main methods). EuSCAPE short read assemblies were screened for containment of these using mash and the FP was taken to be the match with greatest containment. There were no isolates assigned to FPs3/4/5/6 that had linkable phenotypic data.
